# Identification of compounds enhancing *Aqp4* stop codon readthrough

**DOI:** 10.1101/2020.11.01.363945

**Authors:** D. Sapkota, J.D. Dougherty

**Author notes:** **Contact Information:** Dr. Joseph Dougherty, Department of Genetics, Campus Box 8232, 4566 Scott Ave., St. Louis, MO. 63110-1093, E. **Author contribution** D.S. performed most of the experiments, analyzed data, and wrote the paper. J.D.D. acquired the major funding, supervised the project, and helped write the paper.

## Abstract

An unusual stop codon readthrough event generates a conserved C-terminally elongated variant of the water channel protein Aquaporin 4 (AQP4). In the brain, AQP4 is astrocyte-specific, required for normal functioning of the glymphatic system, and involved in the clearance of the Alzheimer’s associated protein Amyloid beta. Further, the readthrough variant is localized exclusively perivascularly, and the perivascular pool of AQP4 is reduced in Alzheimer’s and several other neurological diseases. However, there are currently no means of increasing or restoring the perivascular pool AQP4. Here we identify a compound that can enhance *Aqp4* stop codon readthrough. We screened 2600 compounds, mostly approved drugs and pharmacologically active natural compounds, using a luciferase reporter system. 28 candidate lead compounds were then subjected to a variety of secondary screening steps using orthogonal reporter systems and characterizing dose-response activities. Finally, we tested the top compounds’ abilities to generate readthrough of the endogenous *Aqp4* transcript, identifying Apigenin as an enhancer of this biological phenomenon. This compound can allow modulation of readthrough in experimental systems, mechanistic studies of programmed readthrough, and suggests the potential for modulating Alzheimer’s disease through pharmacological enhancement of perivascular AQP4.

## Introduction

Translation termination determines the length and fate of proteins. This final step of translation is enabled by release factors eRF1 and eRF3 which outcompete amino acyl-tRNAs for the A site of 80S ribosomes that have encountered a stop codon, i.e. UGA, UAG or UAA (Alkalaeva et al., 2006). This results in the hydrolysis of the nascent polypeptide chain from the P site followed by the dissociation of the two ribosomal subunits from the mRNA. A skipped termination, and hence the addition of extra amino acids to the growing polypeptide chain, can result in a protein product that becomes rapidly degraded or acquires untoward functions. It also dissipates cell’s resources since translation is among the most energy-consuming processes. Evolutionarily and thermodynamically, termination is therefore an essential step.

Nevertheless, translation past a stop codon is desirable under certain conditions. Viruses use stop codon readthrough to increase their protein repertoire from a limited amount of genetic material (Firth and Brierley, 2012). Higher-order organisms, from yeasts to humans, also use a programmed readthrough in a subset of transcripts(Dunn et al., 2013; Loughran et al., 2014; Sapkota et al., 2019). Studies on a few of these transcripts have shown that the C-terminally extended protein variants arising from readthrough can gain new properties (Loughran et al., 2018; Schueren et al., 2014).

In the case of *Aqp4*, which encodes an astrocyte-specific water channel protein Aquaporin 4 (AQP4), we and others have recently shown that the readthrough-extended AQP4 is exclusively perivascular in the brain whereas the normal-length AQP4 is restricted away from blood vessels (De Bellis et al., 2017; Sapkota et al., 2019). Importantly, AQP4, specifically its perivascular pool, is a key mediator of the glymphatic system, which clears amyloid beta and other waste products from the brain (Iliff et al., 2012; Mestre et al., 2018). Furthermore, perivascular AQP4 is reduced in Alzheimer’s disease, with the extent of reduction being proportional to amyloid-beta levels (Nilssone et al., 2011; Zeppenfeld et al., 2017). Thus, drugs that promote stop codon readthrough may increase the perivascular pool of AQP4 and hence expedite the removal of amyloid-beta from the brain. Further, such drugs may facilitate studies into the molecular mechanisms by which programmed readthrough occurs.

Although Gentamicin and PTC124 are purported to modulate readthrough of all transcripts, the former does not exert desired effect at subtoxic doses (Linde and Kerem, 2008), and the latter appears to have modulated the Firefly luciferase used in the original screening and hence not to be a genuine modulator at all (Auld et al., 2009; Chowdhury et al., 2018). Therefore, there is a need for additional secondary screens and rigorous validation of hits.

We have previously utilized a dual-luciferase assay to quantify stop codon readthrough of fragments from several brain transcripts (Sapkota et al., 2019). Here, we adapt the same assay to screen compounds for their ability to modulate *Aqp4* stop codon readthrough. We then validate hits using orthogonal approaches independent of luciferase. Finally, we confirm the action of a candidate compound on endogenous *Aqp4* transcript using primary astrocyte cultures.

## Results

### High-throughput screening identifies compounds modulating readthrough of *Aqp4* sequence

The fact that stop codon readthrough generates an elongated AQP4 variant that is required for efficient removal of Amyloid beta suggests that enhancing the phenomenon can be of translational significance. We therefore sought to screen compounds for their ability to enhance *Aqp4* readthrough using a fragment of the transcript we previously demonstrated as supporting readthrough activity (**Fig. 1A)**. We focused on compounds likely to be potential drug candidates surveying a collection of 2560 compounds (Spectrum library), nearly half of which have reached clinical trials in the United States. The remainder are either drugs sold in Asia and Europe or natural products and their derivatives.

**Fig 1.**
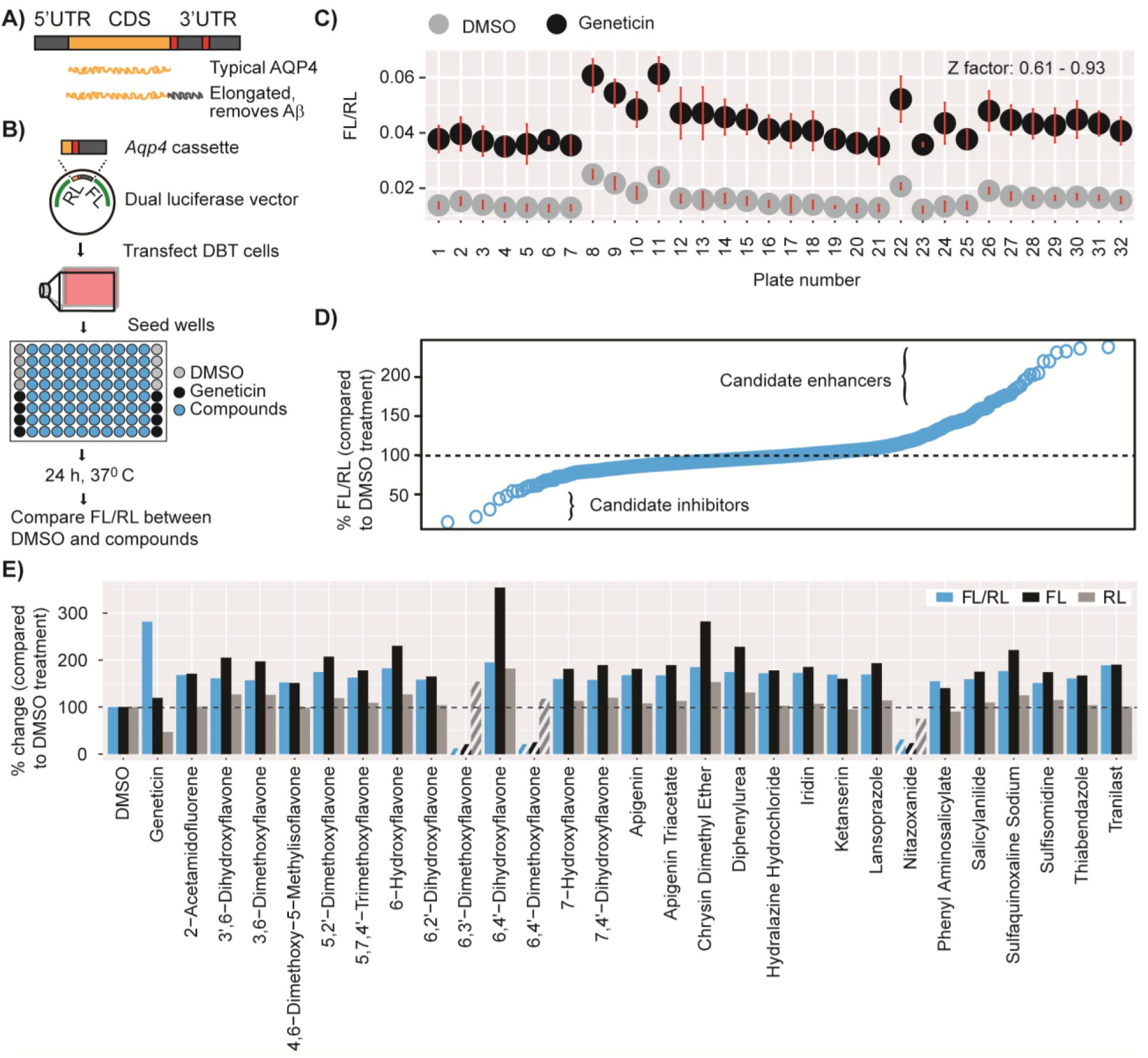
High-throughput screening identifies readthrough modulators. **(A)** Schematic illustration of *Aqp4* readthrough. **(B)** Screening procedure. Transfected cells were seeded into wells pre-spotted with compounds, DMSO (no-treatment control), or Geneticin (positive control). The last column received control cells in which the *Aqp4* stop in the vector was mutated to a sense codon. FL= Firefly luciferase, RL = Renilla luciferase. **(C)** Separation bands between the no-treatment and positive controls and a summary of Z-factors for the plates used in screening. Dots and error bars represent means and 3SD, respectively. **(D)** Quantile analysis of the screening results. **(E)** Compounds enhancing (solid bars) or inhibiting (patterned bars) FL/RL values by at least 50%. The change in FL/RL is largely due to the modulation of FL, not RL.

We began by evaluating a dual luciferase assay that we have previously used to quantify readthrough (Sapkota et al., 2019) for its suitability as a screening approach. This assay utilizes a dual luciferase vector with an *Aqp4* test cassette cloned in between Renilla and Firefly luciferases in such a way that the former is expressed constitutively, but the latter only if the *Aqp4* stop codon is read past by ribosomes. Readthrough rate is then quantified as the ratio of Firefly activity to Renilla activity. We transfected Delayed Brain Tumor (DBT) cells, an astrocyte-like tumor cell line, with this vector and subsequently treated the cells with a series of concentrations of Geneticin, which is an antibiotic known to enhance stop codon readthrough. We found that FL/RL values are highly responsive to Geneticin concentrations (R2 = 0.99) (not shown). Furthermore, the Z-factor, which measures the separation band between the treated cells and untreated cells, was well within the acceptable range of 0.5 and 1 as described (Zhang et al., 1999), thus indicating that the assay is suitable for identifying hits with confidence.

We then screened the Spectrum Library by spotting 96-well plates with 10 μM of each compound, and Geneticin and no-treatment (DMSO) controls; seeding the plates with cells pre-transfected with dual luciferase vector; and measuring FL and RL after 24 h **(Fig. 1B)**. We found that the Z-factor was at least 0.9 for most of the plates and never less than 0.6, demonstrating the robustness of the screening **(Fig. 1C)**. A quartile analysis revealed that most of the compounds did not alter the FL/RL ratio when compared to untreated wells, whereas a few clearly did so **(Fig. 1D)**. Twenty-eight compounds altered the ratio by at least 50%, with 25 increasing and 3 decreasing it **(Fig. 1E)**. Importantly, all twenty-eight hits acted by inducing pronounced changes in FL, which depends on readthrough, but not RL, which is independent of readthrough **(Fig. 1E)**. Therefore, we decided to further validate a set of these hits, which included antiparasitics (thiabendazole, sulphaquinoxaline), an antihypertensive (ketanserin), a common dietary flavonoid (apigenin), and a variety of other flavones.

### Independent assays confirm two compounds as genuine modulators of *Aqp4* readthrough

We started the validation experiments by repeating the dual luciferase assay with 10 μM of the 28 hits in independent cultures. A majority of these compounds caused expected changes in FL/RL, although the magnitudes of changes were somewhat smaller than in the primary screening, possibly due to differences in liquid handling and luciferase detection systems in the two experiments (see methods). As in screening, these changes were largely due to the modulation of FL but not RL (not shown). In total, 11 compounds altered FL/RL by at least 30% when compared to DMSO controls, with 9 enhancing and 2 inhibiting it **(Fig. 2A)**. In parallel, we tested the hypothesis that compounds should cease to exert their effect if FL is already translated at 100%. For this purpose, we mutated the TGA stop codon of the *Aqp4* test cassette cloned in the dual luciferase vector to a sense codon (TGG). We found that all 25 putative enhancers stopped exhibiting their effect **(Fig. 2A)**. The 3 putative inhibitors, however, continued to reduce FL/RL, suggesting they might be impacting translation in general or FL independently of readthrough. They were thus excluded as candidates.

**Fig 2.**
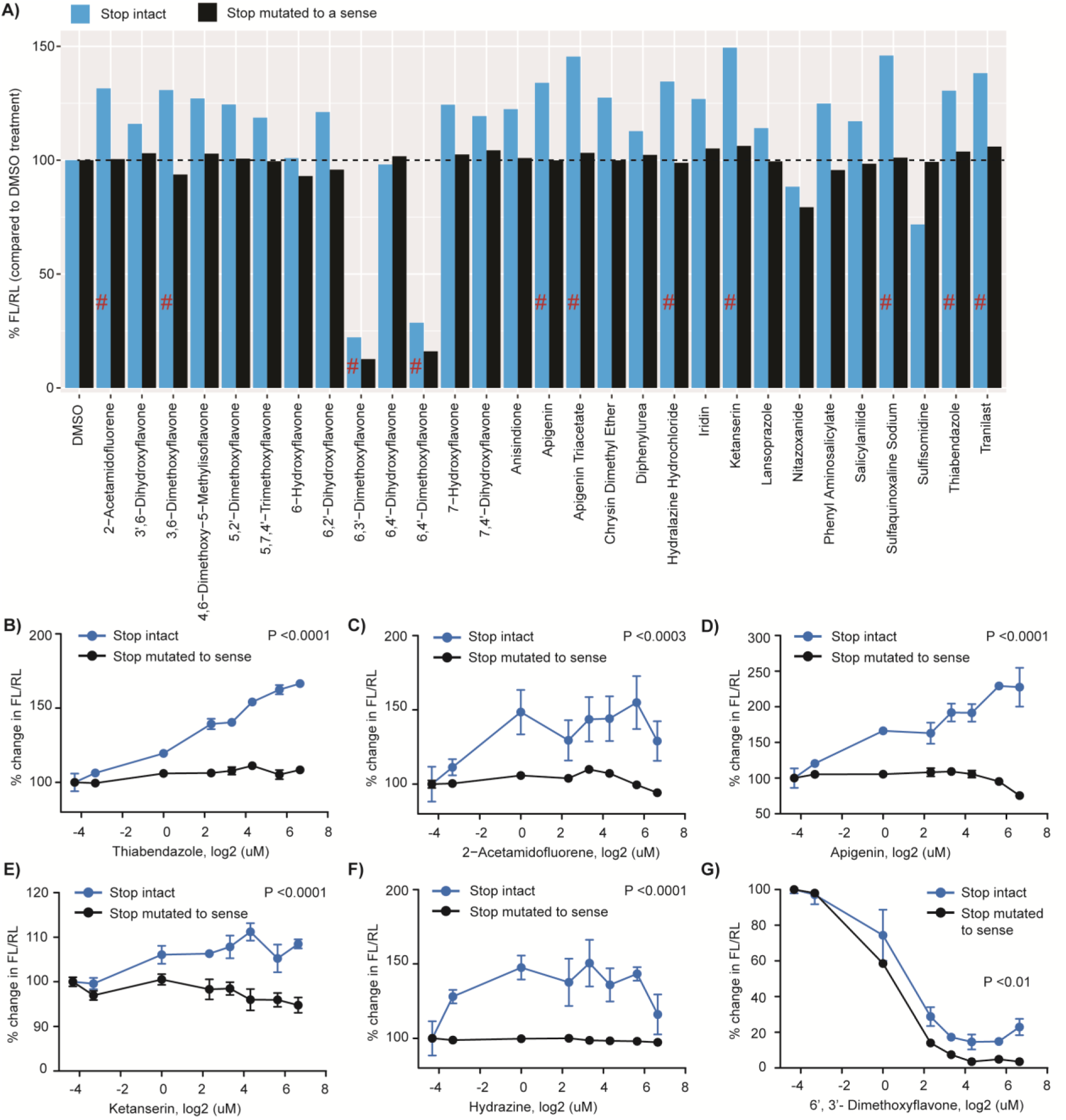
Validation of candidate compounds. **(A)** Retesting with dual-luciferase vectors in which the *Aqp4* stop codon is either intact (blue) or mutated to a sense codon (black). Pound signs indicate compounds altering readthrough by at least 30%. Enhancers cease activity in the absence of stop codon. **(B-G)** Dose-response curves for candidate compounds. Concentrations are log2-transformed. For log transformation of 0 uM (DMSO alone), a value of 0.05 uM was used. Enhancers cease to be active (B-F), whereas an inhibitor continues to be active (G) in the absence of stop codon. Anova followed by a post hoc Tukey was used to compare the means for the two constructs across concentrations.

We also performed a dose-response experiment and, as expected, found all of the tested enhancers act dose-dependently when the test cassette is used but not when the stop-to-sense positive control is used **(Fig. 2B-F)**. In contrast, a tested inhibitor acted dose-dependently with both constructs, again suggesting an impact on overall translation **(Fig. 2G)** or other readthrough-independent mechanisms. These results suggested that the putative enhancers were likely to be genuine modulators of readthrough.

Next, we wanted to further exclude any candidates that altered the FL/RL ratio due to direct activity on the luciferases (i.e. modulating the enzymes rather than actual readthrough). We therefore developed an orthogonal system using a fluorescence-based assay wherein TdTomato/GFP is equivalent to FL/RL **(Fig. 3A)**. Unlike the dual-luciferase assay, however, the fluorescence assay was not sufficiently sensitive for use in DBT cells and required more a more easily transfectable N2a cell line (see methods). Thus, we treated transfected N2a cells with the minimum effective doses of candidate enhancers, as identified from the dose-response experiment, and quantified TdTomato and GFP intensities by plate scanning. We found that two candidate enhancers, Apigenin and Sulphaquinoxaline, significantly increase the TdTomato/GFP ratio when compared to DMSO controls) as expected **(Fig. 3B**. The remaining inhibitor (6’3’ Dimethoxyflavone, 10 μM) did not alter this ratio, indicating it probably inhibits FL directly. We also designed a stop-to-sense positive control fluorescence vector to test whether Apigenin and Sulphaquinoxaline will show activity in a vector that already allows 100% ‘readthrough’ as had been done with the luciferase constructs. Sulphaquinoxaline ceased its activity **(Fig. 3B)**, suggesting it specifically modulates readthrough. Unlike on the luciferase construct, Apigenin showed partial activity on the stop-to-sense construct, **(Fig. 3B)**, indicating while it modulates readthrough it possibly also has a degree of non-specific action on TdTomato or GFP intensity. The activity on this mutated construct, however, was lower than that on the ‘stop intact’ construct; thus, we carried through both compounds to the final step of screening.

**Fig 3.**
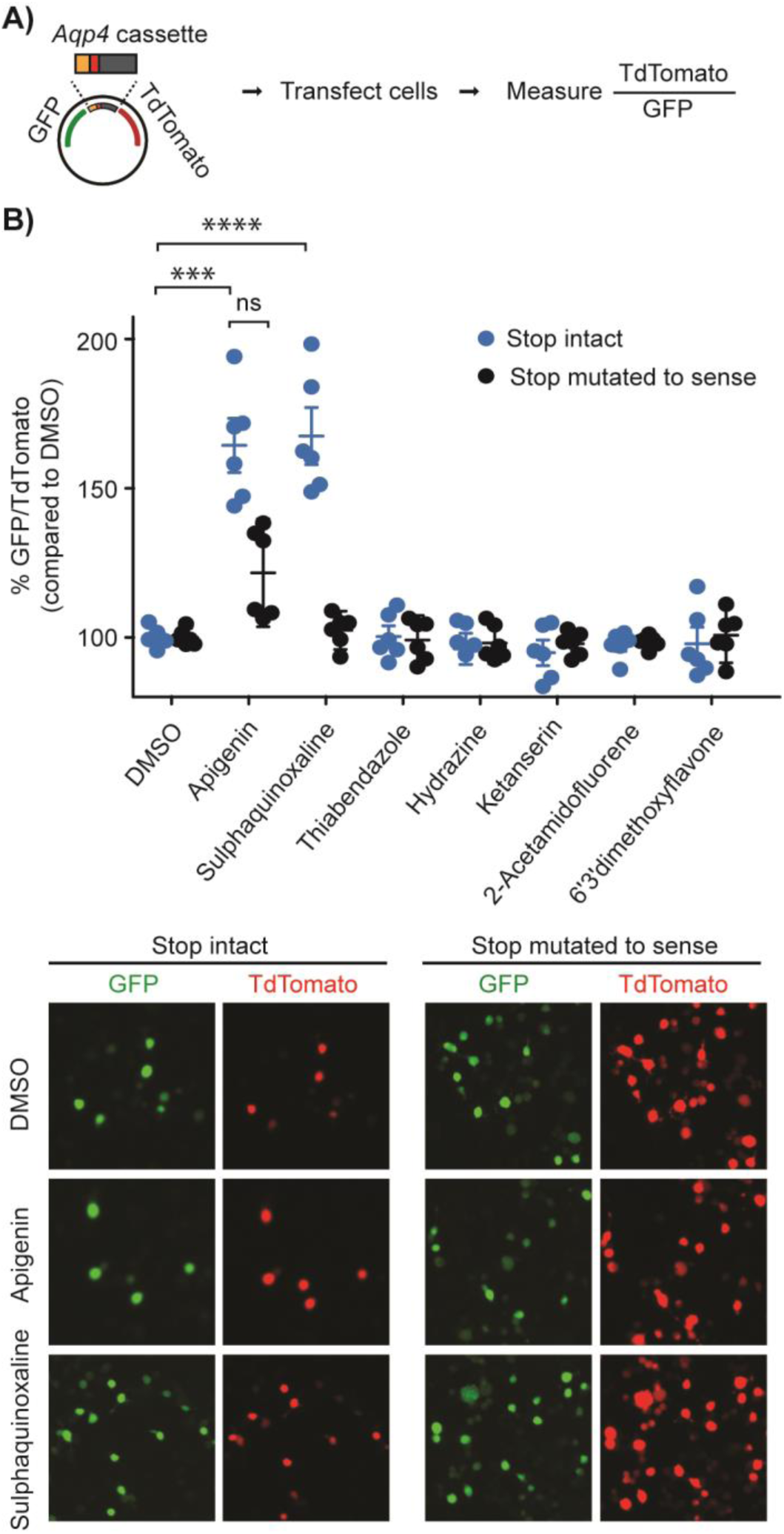
Fluorescence assay validates Apigenin and Sulphaquinoxaline as readthrough enhancers. **(A)** Assay design. TdTomato/GFP is analogous to FL/RL. **(B)** Upper: Assay with dual-fluorescence vectors in which the *Aqp4* stop codon is either intact (blue) or mutated to a sense codon (black). Lower: Representative images for Apigenin and Sulphaquinoxaline. ****, p-value ≤0.0001; ***, p-value ≤0.001, *t*-test.

### Apigenin enhances readthrough of endogenous *Aqp4* transcript

Finally, a *bona fide* modulator should work not just on the cloned fragments of *Aqp4* in our reporters, but also on the endogenous transcript. Thus, we treated primary mouse astrocytes with compounds and quantified the levels of C-terminally extended AQP4 variant, using a dot blot assay with an antibody that specifically recognizes this AQP4 variant (Sapkota et al., 2019). We observed that Apigenin significantly upregulated the expression of extended AQP4, whereas other candidate compounds failed to do so **(Fig. 4A, B)**. These results confirm that Apigenin indeed increases the biosynthesis of readthrough version of endogenous AQP4 in live cells and is a strong candidate for future pre-clinical studies aiming to identify therapies to enhance the perivascular pool of AQP4.

**Fig 4.**
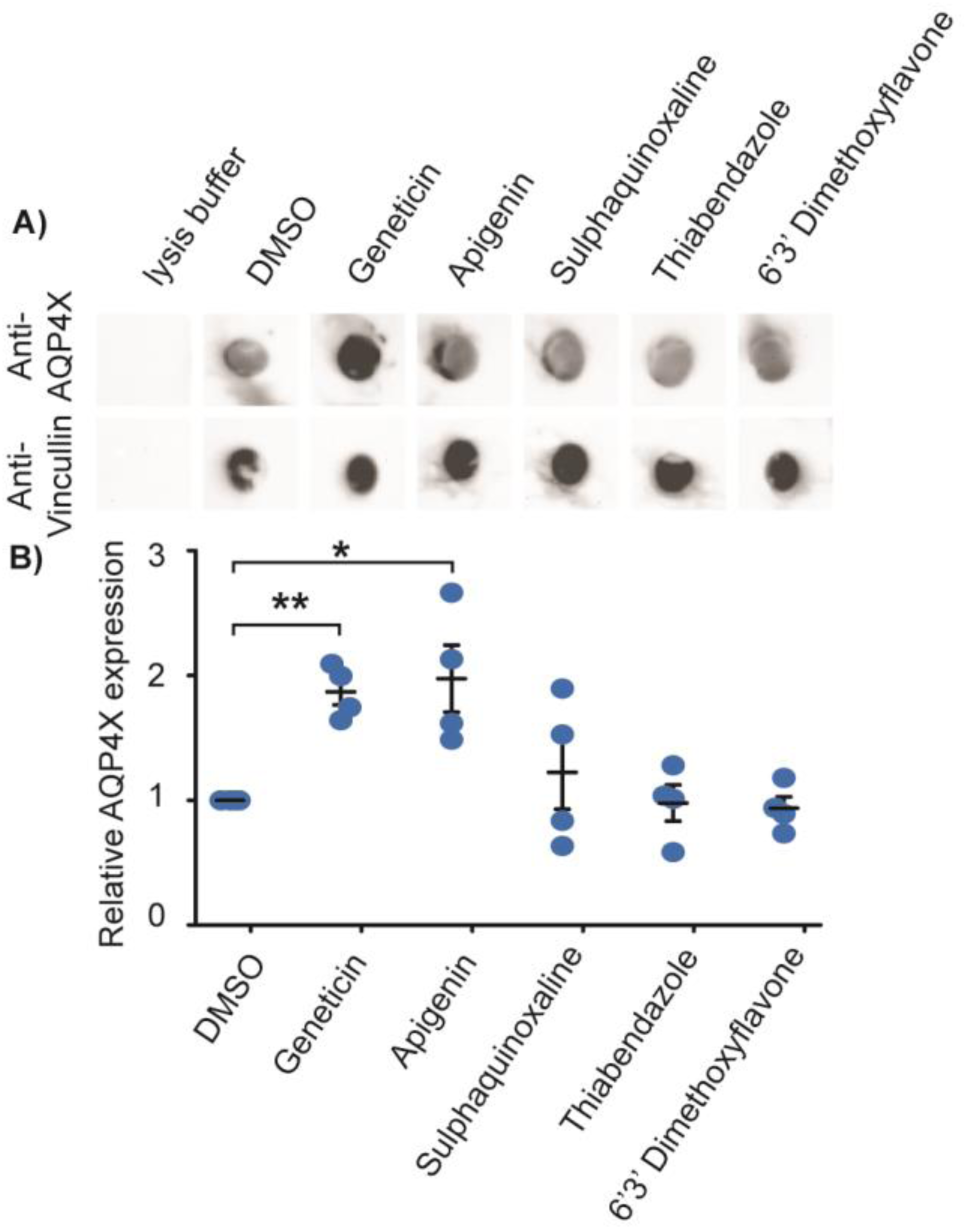
Apigenin enhances readthrough of endogenous *Aqp4* transcript. Representative dot blots for readthrough-extended AQP4 (AQP4X) and Vincullin in primary astrocytes with or without treatments (top). AQP4X expression as normalized to Vincullin (lower). *t*-test comparing DMSO and compounds. **, p-value ≤0.01; *, p-value ≤0.05.

## Discussion and Conclusion

Through a series of screens and counterscreens, two orthogonal reporters, and a readthrough-specific antibody, we have identified Apigenin as a modulator of stop codon readthrough for AQP4 in living cells. This compound may serve as a useful tool for investigating the molecular mechanisms behind programmed stop codon readthrough in *Aqp4*, as well as understanding the biology of enhanced readthrough. Furthermore, it can and has been administered to vertebrates, highlighting potential translational applications. Indeed Apigenin has been shown to reduce plaque burden and improve memory function in a preclinical model of Alzheimer’s disease (Zhao et al., 2013), though any relationship between this compound and stop codon readthrough was not suspected at that time. Revisiting these studies will be essential to determine if Apigenin can efficiently promote Amyloid beta clearance and reduce Alzheimer’s pathology in the brain, and if this depends specifically on its readthrough modulating activity.

Apigenin is a flavonoid (4′,5,7-trihydroxyflavone) that is abundant in edible plants and plant-derived drinks, including parsley, artichoke, mint, spinach, celery, citrus juice, tea and wine (Hostetler et al., 2017). The compound has been found to increase the blood levels of antioxidant enzymes glutathione reductase and superoxide dismutase in humans (Nielsen et al., 1999) and inhibit tumor necrosis factor-α (TNF)-induced proinflammatory gene expression in cell lines (Ruiz and Haller, 2006). Whether Apigenin influences terminating ribosomes, the release factors, or *cis* elements with roles in translation termination, however, remains unknown. Considering our screen and rigorous set of validation tests that Apigenin has passed, the prospect of utilizing this compound in mechanistic studies as well as preclinical studies based on *Aqp4* readthrough is particularly appealing. If it can indeed increase perivascular Aqp4 and thence clearance of Amyloid beta and other waste products in the brain, then it or derived compounds could have potential therapeutic applications in neurodegenerative diseases.

## Materials and Methods

### Cell culture and transfection

Delayed brain tumor (DBT) (Kumanishi et al., 1973) cells were used for high-throughput screening and all other dual luciferase assays. N2a cells (ATCC, CCL-131) were used for fluorescence-based assays. Cells were maintained in Dulbecco’s Modified Eagle Medium (DMEM) supplemented with 100 U/ml each of penicillin and streptomycin and 5% heat inactivated fetal calf serum (Sigma, F4135) and at 37°C in a humidified atmosphere with 5% CO_2_.

Cells were batch-transfected at around 80% confluency in T25 flasks, using Lipofectamine LTX reagents (Invitrogen) and 7.5 μg plasmid, and following manufacturer’s instructions. After 6 h, cells were detached and used for screening or validation experiments.

### Vectors for readthrough assay

For luciferase-based assays, an *Aqp4* test cassette (the last 15 nts of the coding sequence + annotated stop codon + 84 nts before the 2nd stop) was cloned between the XhoI and BglII sites of pdLUC, a dual-luciferase vector, such that Renilla luciferase would be expressed constitutively but Firefly luciferase would be expressed only if the stop codon was read by ribosomes (Fixsen and Howard, 2010). Readthrough rate was quantified as the ratio of Firefly activity to Renilla activity in lysates from cells expressing the test vector. The annotated stop codon (TGA) was mutated to TGG in a positive control vector to allow 100% readthrough.

Similarly, for fluorescence-based assays, the test and control cassettes were cloned between GFP and TdTomato using MreI and RsrII sites in an FCIV vector backbone (Hope Center, Washington University). The readthrough rate was quantified as TdTomato intensity/GFP intensity.

### High-throughput screening

A Spectrum collection of 2560 compounds (10 mM in DMSO) was purchased from MicroSource Discovery Systems, Inc. USA. 100 nL of compounds were dispensed in individual wells of white-wall clear-bottom 96-well plates (Corning, 3903), leaving the first and last columns. 1 μL of 10% DMSO was dispensed in the upper four wells of the first and last columns (no treatment control). 1 uL each of 10% DMSO and 50 mg/mL Geneticin, an antibiotic known to enhance stop codon readthrough, (G8168, Sigma), was dispensed in the lower four wells of the two columns (positive control). The pre-spotted wells were seeded with DBT cells harvested from T25 flasks after 6 h of transfection. Wells with compounds and wells of the first column received cells expressing the test *Aqp4* cassette. Wells of the last column received cells expressing the positive control *Aqp4* cassette. Each well received 8 × 10^4^ cells. The final volume in each well was 100 uL, and thus the final concentrations of the compounds, DMSO, and Geneticin were 10 μM, 0.1%, and 500 μg/mL, respectively. Importantly, we pre-confirmed that 0.1% DMSO alone does not impact the readthrough rate.

Plates were incubated for 24 hours at 37°C and assayed for FL and RL using the Dual-Luciferase Assay System and following manufacturer’s instructions (E1980, Promega). Liquid handling and luciferase assay were performed using the integrated automated screening platform (Beckman Coulter) at the High Throughput Screening Center at Washington University. Screening quality was assured by calculating the Z-factor for individual plates using the FL/RL values from the first column as follows and ensuring that it was between 0.5 and 1 as described (Zhang et al., 1999).

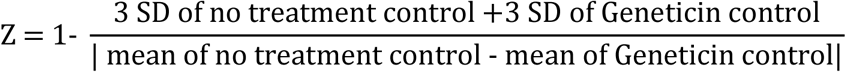

A few compounds induced noticeable cytotoxicity, altered RL values by more than 25% when compared to no treatment control or resulted in FL values close to zero and were excluded from further analysis. Compounds surviving these criteria and altering the FL/RL ratio by at least 50% were considered for further validation experiments.

### Validation using dual-luciferase and fluorescence assays

Dual-luciferase assay-based validation, dose-response, and experiments followed the same overall procedure as used for initial screening, except that Cytation5 with dual injectors (BioTek) was used for dual-luciferase assays and candidate compounds were purchased from Sigma (Sulphaquinoxaline, S7382; Thiabendazole, T5535; Ketanserin, S006; 6’3’-Dimethoxyflavone, CDS007225; Hydrazine 216194; Apigenin, 178278; Tranilast 616400; 2-Acetamidofluorene, A7015).

Fluorescence-based assay also followed the same procedure as described for screening, except that after treatment, individual wells were scanned from the bottom and the mean fluorescence intensities of TdTomato and GFP were quantified using Cytation5. Prior to fluorescence quantitation, the medium was replaced with PBS. Black-walled clear-bottom 96 well plates were used (Corning, 3603). To make this assay consistent with dual-luciferase assay, we first used DBT cells. However, these cells had only around 15% transfection rate and showed TdTomato/GFP ratios that were indistinguishable from the background even after treatment with Geneticin, which is known to enhance readthrough by several folds. N2a cells had around 90% transfection rate and showed TdTomato/GFP ratios highly responsive to Geneticin. Thus, this assay utilized N2a cells. The following minimum effective doses as determined from a dose-response experiment were used: Thiabendazole (20 μM), Apigenin (20 μM), Hydrazine (1 μM), Ketanserin (15 μM), 2-Acetamidoflurene (1 μM), and Sulphaquinoxaline (10 μM).

Validation experiments included at least three each of experimental and technical replicates. Statistical tests are mentioned in figure legends.

### Primary astrocyte culture and drug treatment

Four postnatal day 2 mouse pups were anaesthetized in ice, sprayed with 70% ethanol and rapidly decapitated. The brains were harvested, and the cortices dissected out in ice-cold PBS. After removing the meninges, the cortical hemispheres were minced in 2 ml of prewarmed DMEM taken in a 35-mm dish. Trypsin was added to the final concentration of 0.25% and the dish was shaken for 30 min at 37°C. Tissue pieces were triturated in 5 mL DMEM and in the presence of DNAseI in a 15 mL conical tube. Fetal bovine serum was added to the final concentration of 10%, larger tissue pieces were allowed to settle, and the supernatant was centrifuged at 200 xg for 5 min. The pellet was resuspended in complete medium (DMEM + 10% fetal bovine serum + antibiotics) and 1 × 10^6^ cells were plated in T25 flasks. Upon confluency after a few days, the flasks were shaken for 30 min to stir up neurons and oligos, medium was discarded, and the cells were passaged into 6-well culture plates. Confluent growths in 6-well plates were treated for 24 h with 0.1% DMSO, Geneticin (50 mg/mL), Apigenin (20 μM), Sulphaquinoxaline (10 μM), Thiabendazole (20 μM), and 6’3’ Dimethoxyflavone (10 μM) and subjected to dot blot assays.

### Dot blot assay

Cells were washed with ice-cold PBS and lysed in RIPA buffer (50 mM pH 8.0 Tris-HCl, 150 mM NaCl, 1% NP-40, 0.5% sodium deoxycholate, 0.1% SDS, 1mM NaVO4, 1mM NaF, Roche complete protease inhibitor tablet). Lysates were cleared by centrifuging at 16,000 Xg at 4°C for 10 m. Protein concentration was measured using a BCA kit (ThermoScientific). 5 ug of total protein in a volume of 4 uL was loaded onto polyvinyl membrane strips laid on top of a moist filter paper. The blots were air-dried for 1 h, blocked in 5% non-fat dried milk in tris-buffered saline with 0.1% Tween-20 (TBST) for 1 h at room temperature, incubated with anti-AQP4X (1:4000, Cell Signaling Technology, #60789) or anti-Vincullin (Abcam, #ab129002) in block overnight at 4°C, washed three times with TBST, incubated with HRP-conjugated secondary antibody in block (1:5000, BioRad) for 1 h at room temperature, washed three times with TBST, and finally developed using ECL reagents (BioRad) and My ECL imager (ThermoScientific).

## Acknowledgements

We thank Maxene Ilagan at the High-Throughput Screening Center at Washington University in St. Louis for her technical support with screening, Chengran Yang for help with cloning, Gary Loughran for the dual-luciferase plasmid, and members of the J.D.D. laboratory for scientific discussions. This work was supported by the NIH (1R01NS102272) and Washington University Hope Center to J.D.D. and NIA (K99AG061231) to DS.

